# Influence of Supraphysiologic Biomaterial Stiffness on Ventricular Mechanics and Myocardial Infarct Reinforcement

**DOI:** 10.1101/2020.05.20.107276

**Authors:** Ravi K. Ghanta, Yunge Zhao, Aarthi Pugazenthi, Mathew J. Wall, Lauren N. Russell, Kyle J. Lampe

## Abstract

Injectable intramyocardial biomaterials have promise to limit adverse ventricular remodeling through mechanical and biologic mechanisms. While some success has been observed by injecting materials to regenerate new tissue, optimal biomaterial stiffness to thicken and stiffen infarcted myocardium to limit adverse remodeling has not been determined. In this work, we present an in-vivo study of the impact of biomaterial stiffness over a wide range of stiffness moduli on ventricular mechanics. We utilized injectable methacrylated polyethylene glycol (PEG) hydrogels fabricated at 3 different mechanical moduli: 5 kPa (low), 25 kPa (medium/myocardium), and 250 kPa (high/supraphysiologic). We demonstrate that the supraphysiological high stiffness favorably alters post-infarct ventricular mechanics and prevents negative tissue remodeling. Lower stiffness materials do not alter mechanics and thus to be effective, must instead target biological reparative mechanisms. These results may influence rationale design criteria for biomaterials developed for infarct reinforcement therapy.

## INTRODUCTION

Acellular intramyocardial biomaterials have shown promise for infarct reinforcement to prevent adverse left ventricular (LV) remodeling post-myocardial infarction (MI).[1-3] Post-infarct remodeling is a maladaptive, progressive process characterized by LV chamber dilation, wall thinning, and infarct expansion. Early infarct expansion and infarct borderzone expansion leads to a poor long-term prognosis and development of heart failure.[4, 5] Post-infarction, infarct areas are initially compliant and dyskinetic allowing for infarct expansion. Consequently, therapies of infarct restraint have been proposed to prevent infarct expansion and adverse ventricular remodeling.[6-8]

Intramyocardial biomaterials were initially hypothesized to prevent ventricular remodeling through mechanical mechanisms, namely augmentation of wall thickness, infarct stiffening, and reduction of mechanical wall stress.[9, 10] A variety of synthetic and natural biomaterials, with a wide range of stiffness and biologic properties, have been developed for infarct reinforcement in animal models and patient clinical trials. [2, 3, 10, 11] Virtually all prior studies have utilized naturally derived (hyaluronic acid, alginate, extra-cellular matrix) biomaterials, which may be costly to derive and difficult to modify. It is also unclear if the functional benefit seen in prior studies with these relatively soft materials was the result of a mechanical effect or a response to a bioactive component of the material. [10, 12] Rane and colleagues found no benefit for soft (0.5 kPA) bioinert poly-ethylene glycol (PEG) injection in a small animal model.[10] Computer simulations and finite element analyses have demonstrated that biomaterial stiffness, degradation, and injection location can significantly alter efficacy of therapy.[9, 13, 14] Matsumura and colleagues recently were the first to demonstrate efficacy of a synthetic biomaterial, utilizing a relatively stiff (275kPA) material.[15] Synthetic materials have numerous advantages, including the ability to tune material degradation and stiffness properties. Few studies have evaluated the effect of biomaterial stiffness on heart function *in-vivo.* These prior studies have evaluated a limited range of stiffness and have not evaluated supraphysiologic stiff materials. Consequently, rational design criteria for biomaterials and mechanical dosing are not established.

The aim of this study is to determine the role of intramyocardial biomaterial stiffness on regional post-infarct ventricular mechanics and determine the optimal stiffness levels to achieve efficacy. We hypothesize that biomaterial stiffness greater than normal myocardium is optimal for improving efficacy of intra-myocardial acellular injection therapy. To evaluate this hypothesis, injectable methacrylated polyethylene glycol (PEG) hydrogels capable of wide range of stiffness properties were utilized to determine the acute and chronic effects of injection on regional ventricular mechanics and stiffness.

## MATERIALS & METHODS

### Study Design Overview

This study was divided into 2 parts. In part 1, the acute effect of infarct reinforcement on global and regional ventricular mechanics in a rat left anterior descending (LAD) ligation model was evaluated. Three different mechanical doses of PEG hydrogel plus control (saline) were utilized. PEG hydrogels were fabricated using reduction/oxidation polymerization at 3 different mechanical doses: 4.9 +/- 0.3 kPa (Low stiffness; similar to most prior injectable materials), 24.6 +/- 1.0 kPa (Normal stiffness; similar to normal rat myocardium), and 249 +/- 74 kPa (Supraphysiologic stiffness).[16] Hydrogels were fabricated according to previously described methods and stiffness was confirmed with rheometric measurement.[17] For each mechanical dose, the global and regional ventricular mechanics at baseline, 30 minutes post-infarction, and 30 minutes post-infarct stiffening were measured and compared. Pressure-volume analysis was then performed for each subject (n=20; 5 per group) at 3 time points. In Part 2, the chronic effect of infarct stiffening on ventricular function and remodeling was evaluated using cardiac MRI. Animals underwent LAD ligation followed by immediate injection of either control (saline), low mechanical dose (5kPA) PEG or supraphysiologic high mechanical dose (250kPa) PEG. Animals were recovered from surgery and cardiac MRI was performed 4 weeks post-injection (n=9; 3 per group). We then evaluated LV remodeling and regional strain. All animals were treated and cared for in compliance with the Guide for Care and Use of Laboratory Animals published by the National Institutes of Health (NIH publication 86-23, revised 1996). The protocol was approved by the Institutional Animal Care and Use Committees at the University of Virginia and Baylor College of Medicine. For this study, 8 week old male Sprague Dawley rats with a mean weight of 300 ± 15gms were utilized.

### PEG Hydrogel Preparation and Characterization

4600 or 1000 g/mol molecular weight (MW) functionalized PEG were used in the formulation of the hydrogels. 4600 MW PEG (Millipore Sigma, St. Louis, Missouri) was methacrylated to ∼80% functionalization according to previous protocols.[17] 1000 MW PEG diacrylate was purchased from Sigma with a manufacturer-reported functionalization of 70%. Hydrogel solutions consisted of 12.5% 4600 MW PEG dimethacrylate, 30% 4600 MW PEG dimethacrylate, or 30% 1000 MW PEG diacrylate to create hydrogels with storage moduli of approximately 5, 25, and 250 kPa, respectively. For each hydrogel formulation, reducing and oxidizing solutions were made based on prior protocols.[18, 19] The reducing solution consisted of methacrylate or diacrylate PEG (4600 or 1000 MW, respectively), ferrous gluconate dihydrate (Sigma), and ascorbic acid (Millipore Sigma, St. Louis, Missouri) in water; the oxidizing solution was functionalized PEG and tertbutyl hydroperoxide (Millipore Sigma, St. Louis, Missouri) in water (**Figure 1**). Ferrous gluconate dehydrate and tertbutyl hydroperoxide were added at 0.005:1 mol ratios relative to PEG in reducing and oxidizing solutions, respectively. Ascorbic acid was added relative to the ferrous gluconate dehydrate at a 3.15:1 mole ratio to inhibit spontaneous polymerization[19]. Hydrogel plateau storage moduli were determined from oscillatory shear rheology on an Anton Paar MCR 301 rheometer with a fixed frequency of 10 Hz (n=6).

**Figure 1.**
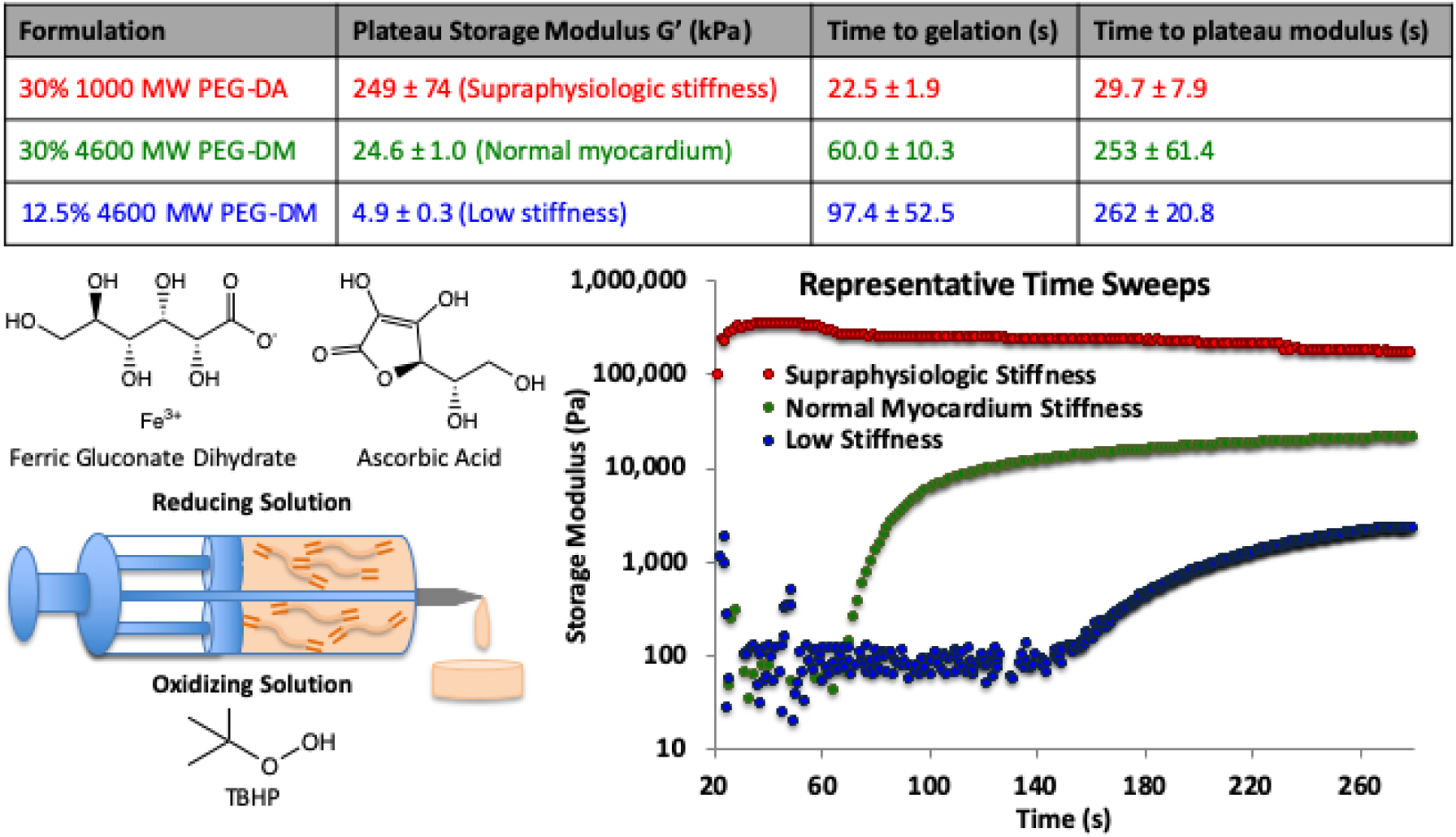
Rheometric characteristics of PEG formulations. Crosslinkable PEG formulations were developed to create hydrogels that approximated normal myocardium and values an order of magnitude lower and greater. Hydrogels were created by varying PEG molecular weight and weight percent in solution during crosslinking, which was performed by mixing reducing solutions of functionalized PEG, ferric gluconate dehydrate, and ascorbic acid, with the oxidizing solution of functionalized PEG and TBHP. Time to gelation was less than 2 minutes in all cases and the time to plateau modulus less than 5 minutes. The formulations yielded hydrogels with plateau moduli spanning two three orders of magnitude. 4.9 (low), 24.6 (normal myocardium), and 249 (supraphysiologic) kPa gels were used to create low, medium, and high mechanical doses, respectively, in infarcted rat hearts.

### Acute Rat Infarction and Instrumentation

For Part 1 of this study, the effect of infarct stiffening on regional ventricular mechanics was evaluated. Animals were anesthetized with inhalational isoflurane in 100% O_2_ (5.0% induction; 2.5% maintenance). Endotracheal intubation was performed and the animals ventilated with positive-pressure ventilation. A median sternotomy was then performed. The pericardium and pleural spaces were opened. The inferior vena cava (IVC) was encircled with a silk suture to allow for caval occlusions. Six sonomicrometer crystals (Sonometrics, London, Ontario, Canada) were sewn to the epicardial surface of the LV as illustrated in **Figure 2**. Crystals were placed on the LV at the base, apex, anterior/septum, inferior/posterior wall, lateral wall, and 2-4mm left of the LAD at the level of the planned ligation. A Millar SP-671 pressure transducer (Millar Instruments, Houston, Texas) was inserted in the LV cavity through the apex. Data were acquired at 312 Hz by using the commercial software SonoLab (Sonometrics). Baseline recordings of LV pressure and crystal pairs were performed. A caval occlusion was performed while recording hemodynamic data to allow measurements over a range of preload and afterload conditions. The proximal to mid LAD was then ligated with a 7-0 poly-propylene suture (Ethicon, Somerville, New Jersey). Infarct was noted visually and then confirmed by sonomicrometry recordings. If an insufficient infarct was identified, additional sutures were placed. Following 30 minutes, hemodynamic recordings were obtained of the post-infarction state. The mortality of ligation was 55%. Survivors were divided into 4 groups for infarct reinforcement: control, low PEG stiffness (5kPA), medium PEG stiffness (25kPA), and high PEG stiffness (250kPA). PEG or control was injected at 4 sites corresponding to 4 quadrants of the infarct. A total of 0.2mL of saline or PEG was injected into the LV wall, which led to a visual change in the LV wall confirming delivery. No animals died secondary to injection. To minimize leakage of biomaterial, needles were inserted at a 45 degree angle to the heart surface and slowly withdrawn with direct compression of a cotton tip applicator which was held over the exit site for 3 minutes. Hemodynamic recordings were obtained 30 minutes post injection. Animals were then euthanized.

**Figure 2.**
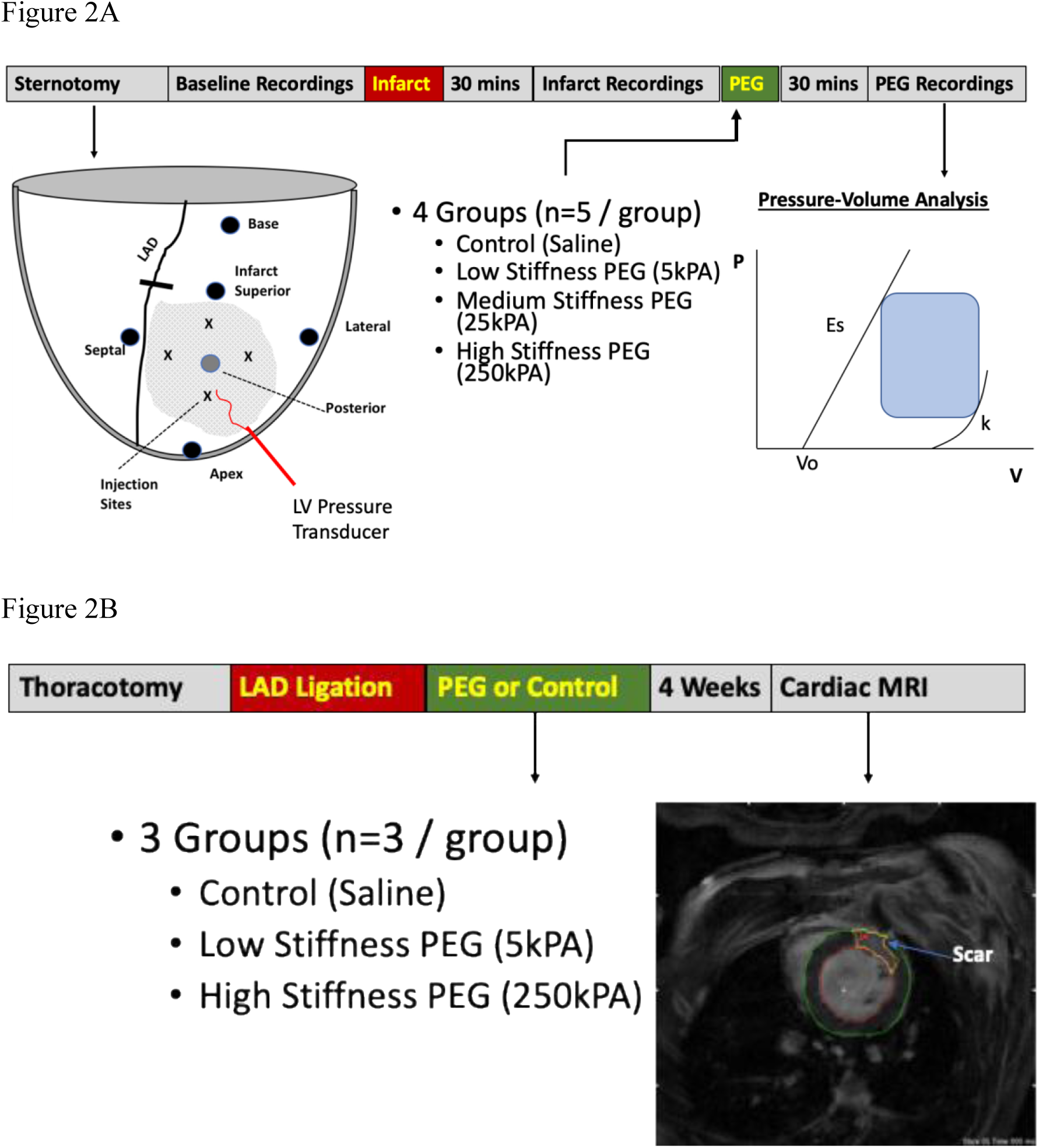
Overview of experimental protocol for (A) acute study and (B) chronic study. For the acute study sonomicrometer crystals were placed on the anterior, lateral, and apex (black circles) and the posterior surface of the heart (grey circle). Pressure transducer placed via the apex to measure LV pressure. LAD Ligation performed (black bar) to induce large anterior wall infarction.

### Sonomicrometry and Pressure Volume Analysis

Sonomicrometry and hemodynamic data were analyzed using custom designed algorithms with MATLAB (The Mathworks, Natick MA) and SonoLab software (Sonometrics).[6, 20] The sonomicrometry crystal pairs were utilized to determine regional strain and LV volume.[21, 22] **Figure 2** illustrates the crystal locations and pairing utilized to determine regional ventricular function. The base-apex pair constituted the long-axis measurement and the anterior-posterior for short-axis measurement. LV volume was calculated utilizing the base-apex and anterior-posterior segment lengths assuming a truncated ellipsoidal geometry for the LV.[20] The anterior and lateral pair constituted the infarct area circumferential strain. The infarct superior-apex crystal pair constituted the infarct longitudinal strain. Infarct area strain was defined as anterior-lateral * base-apex lengths. Remote area was defined as the triangular region from the apex-posterior, apex-lateral, and lateral-posterior crystal pairs. Remote area strain was defined as the area of these 3 crystal pairs with lateral-posterior set as the base of the triangular region. For each cardiac cycle, end-diastole (ED) was defined as the time immediately before the sharp increase in LV pressure. End-systole (ES) was defined as the point of maximum elastance. The end-systolic pressure volume relationship (ESPVR) was then calculated from the caval occlusion data, by the procedure of Kono et al.[23] We identified the slope of the ESPVR, the systolic elastance (E_s_) and the volume access intercept (V_o_), both well-established measures of global ventricular function. The end-diastolic pressure volume relationship (EDVPR) was modeled exponentially (P=A*exp(*k*\*v), with *k* representing the diastolic stiffness constant.[24] Indices of myocardial oxygen consumption (M_v_O_2_) were then calculated. The tension-time index (TTI) was calculated as the integral of LV pressure with respect to time over the cardiac cycle. We determined the pressure-volume area (PVA), defined as the area under the ESPVR and pressure-volume loop. Stroke work was defined as the area circumscribed by the pressure-volume loop. Mechanical efficiency (ratio of stroke work to PVA) was calculated. Systolic strain was calculated from systolic area minus end-diastolic area. To facilitate comparison, recordings were normalized to end-diastole area. Peak systolic strain was identified as the maximum absolute value of strain throughout systole.

### Chronic Rat Infarction

For the chronic rat infarction model, general anesthesia was induced as described for acute experiments and subjects were endotracheally intubated. A left lateral thoracotomy was performed. The LAD was identified and ligated with a 7-0 poly-propylene suture at the proximal one third level. Visual confirmation of infarction was noted by edema and discoloration of the anterior LV wall. After stability (no fibrillation or agonal contraction) of the animals were confirmed, animals were divided into three groups: control (saline), low stiffness (5KPA), or high stiffness (250kPA) PEG injection. Saline or PEG was injected at 4 sites corresponding to the 4 quadrants of the infarct area for a total injection volume of 0.2mL. No animals died after injection. The thoracotomy was closed and the animal was allowed to recover.

### Cardiac MRI and Analysis

Cardiac MRI was performed at 4 weeks following LAD infarction. A 7 Tesla ClinScan (Bruker, Ettlingen, Germany) was utilized and cines short axis images through the entire left ventricle was performed. All image analysis for global LV size and function was quantified using the freely available software Segment version 3.0 (http://segment.heiberg.se).[25] Presence of infarct was confirmed with inversion recovery gradient echo (IR-GRE) pulse sequence, which allows for hyperenhancement of the infarcted area. Infarct area and wall thickness was quantified using the Segment software. Diastolic function was estimated by the peak first derivative of LV volume over time (LV dV/dt) during diastole.[26-28]

### Statistical Analysis

Results are presented as mean ± standard deviation. Normality was examined with the Shapiro-Wilk test. When there was a normal distribution, means between groups were compared with the paired and unpaired Students’ t-test. Repeated measures data were assessed using a multi-sample repeated measures ANOVA with a post-hoc paired Student’s t test and Holm-Bonferroni correction. When there was not a normal distribution, a Kruskal-Wallis test was performed with a post hoc Wilcoxon signed rank test. A *p*<0.05 was considered to indicate significance. Statistics were performed utilizing STATA version 15 (STAT Corp, College Station, Texas, USA).

## RESULTS

### Hydrogel Properties

Plateau storage moduli were measured at 4.9 +/- 0.3 kPa (Low stiffness; similar to most prior injectable materials), 24.6 +/- 1.0 kPa (Medium stiffness; similar to normal rat myocardium), and 249 +/- 74 kPa (Supraphysiologic stiffness) for the 12.5% 4600 MW PEG dimethacrylate, 30% 4600 MW PEG dimethacrylate, and 30% 1000 MW PEG diacrylate hydrogels, respectively. **Figure 1** illustrates rheometry findings. On the rheometer stage, hydrogel gelation occurred in roughly thirty seconds and reached the plateau storage modulus in two minutes.

### Rat Infarction Model

**Table 1** and **Figure 3A** demonstrate that acute ischemic heart failure was achieved following LAD ligation. Thirty minutes following infarction, animals developed a rightward shift in the ESPVR (V_o_ from 39± 21mL to 233 ± 87mL; p<0.05), increased LV EDV (254 ± 133 to 453 ± 167mL; p<0.05), increased LV EDP (7 ± 1mmHg to 12 ± 2mmHg; p<0.05), and decreased EF (58 ± 13% to 26 ± 4%; p<0.05). Similarly, cardiac energetics were adversely affected 30 minutes following infarction with increased PVA (5.6 ± 1.2 to 7.9 ± 2.3; p<0.05) and decreased mechanical efficiency (79 ± 11% to 60 ± 13%; p<0.05). There was no change in systolic elastance or diastolic stiffness.

**Table 1.** Global Hemodynamics, Mechanics and Energetics at Baseline vs Post-Infarction.

|  | Baseline | Post-Infarction |
| --- | --- | --- |
| **Pressure-Volume** |  |  |
| LV EF, % | 58 ± 13 | 26 ± 4* |
| LV EDV, µL | 254 ± 133 | 453 ± 167* |
| LV EDP, mmHg | 7 ± 1 | 12 ± 3* |
| Ees mmHg/mL | 575 ± 193 | 532 ± 133 |
| Volume Axis Intercept, µL | 39 ± 21 | 233 ± 87* |
| Diastolic Stiffness, mL<sup>-1</sup> | 6 ± 2 | 5 ± 3 |
| **Energetics** |  |  |
| Tension Time Index, mmHg S | 5.5 ± 0.9 | 5.9 ± 0.7 |
| Pressure Volume Area, mmHg mL | 5.6 ± 1.2 | 7.9 ± 2.3* |
| Mechanical Efficiency, % | 79 ± 11 | 60 ± 13* |

EDV, End-Diastolic Volume; EDP, End Diastolic Pressure; EF, Ejection Fraction; Ees, End-Systolic Elastance; \*p<0.05 for paired t-test.

**Figure 3.**
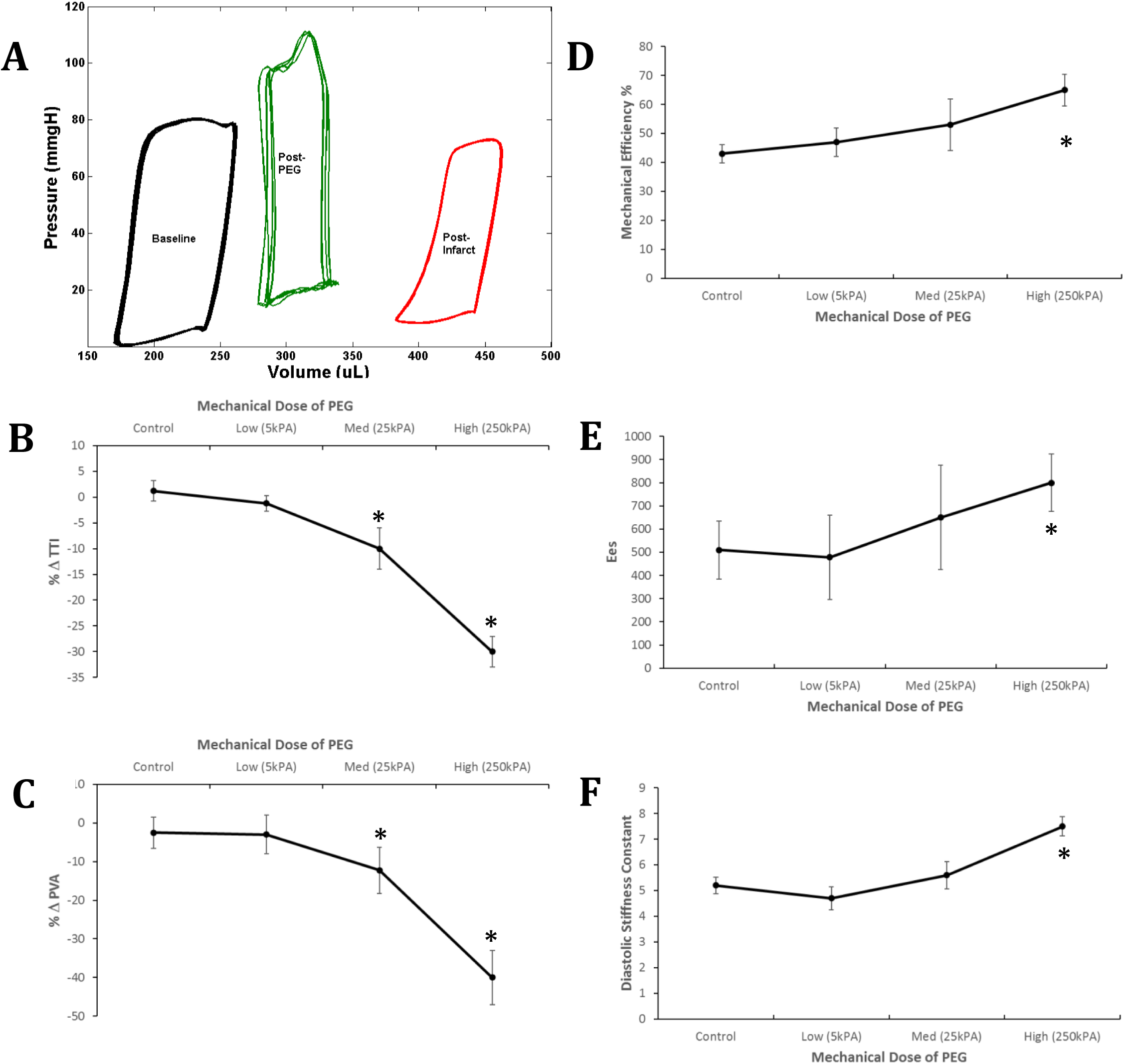
Pressure-Volume Analysis. A) PV loops at baseline, post-infarct, and post-injection of high mechanical dose PEG. Infarction shifts the PV loop rightward. Injection leads to shift of the PV loop leftward. B) TTI decreases with PEG injection relative to post-infarction values. Significant reduction is noted at medium and high mechanical doses of PEG. C) PVA also decreases with PEG injection relative to post-infarction value. Significant reduction is noted at medium and high mechanical PEG doses. D) Mechanical efficiency (stroke work/PVA) improves with high mechanical dose of PEG. E) Systolic elastance increases with high mechanical dose of PEG. F) Diastolic stiffness increases with high mechanical dose of PEG. n=5 subjects/group; *p<0.05 relative to post-infarct.

### Global Ventricular Mechanics and Energetics

At medium and supraphysiologic mechanical doses, infarct stiffening with PEG hydrogels improved global ventricular mechanics and reduced indices of myocardial oxygen consumption (**Figures 3 and 4**). Furthermore, the efficacy was dependent on the mechanical dose of PEG administered. Indices of myocardial oxygen consumption were unchanged with control and low stiffness (5kPA) PEG gels, but decreased after medium stiffness (25kPA) and supraphysiologic stiffness (250kPA) PEG gels were administered. TTI decreased 10 ± 4% (p=0.04) and 30 ± 3% (p<0.01) respectively for medium and supraphysiologic stiffness PEG conditions. Similarly, PVA decreased 12±6% (p=0.03) and 40±7% (p<0.01) respectively for medium and supraphysiologic stiffness PEG gels. Mechanical efficiency improved only with supraphysiologic stiffness PEG gel (50±15% to 75±6%; p<0.01). Systolic elastance only improved with supraphysiologic stiffness PEG gel (from 590 ± 128 to 800 ± 125; p=0.04). Diastolic stiffness constant also increased only with supraphysiologic stiffness PEG gel (from 5.2 ± 0.32 to 7.5± 0.38; p=0.03). All comparisons are pre-injection/post-infarct versus post-injection foreach group.

**Figure 4.**
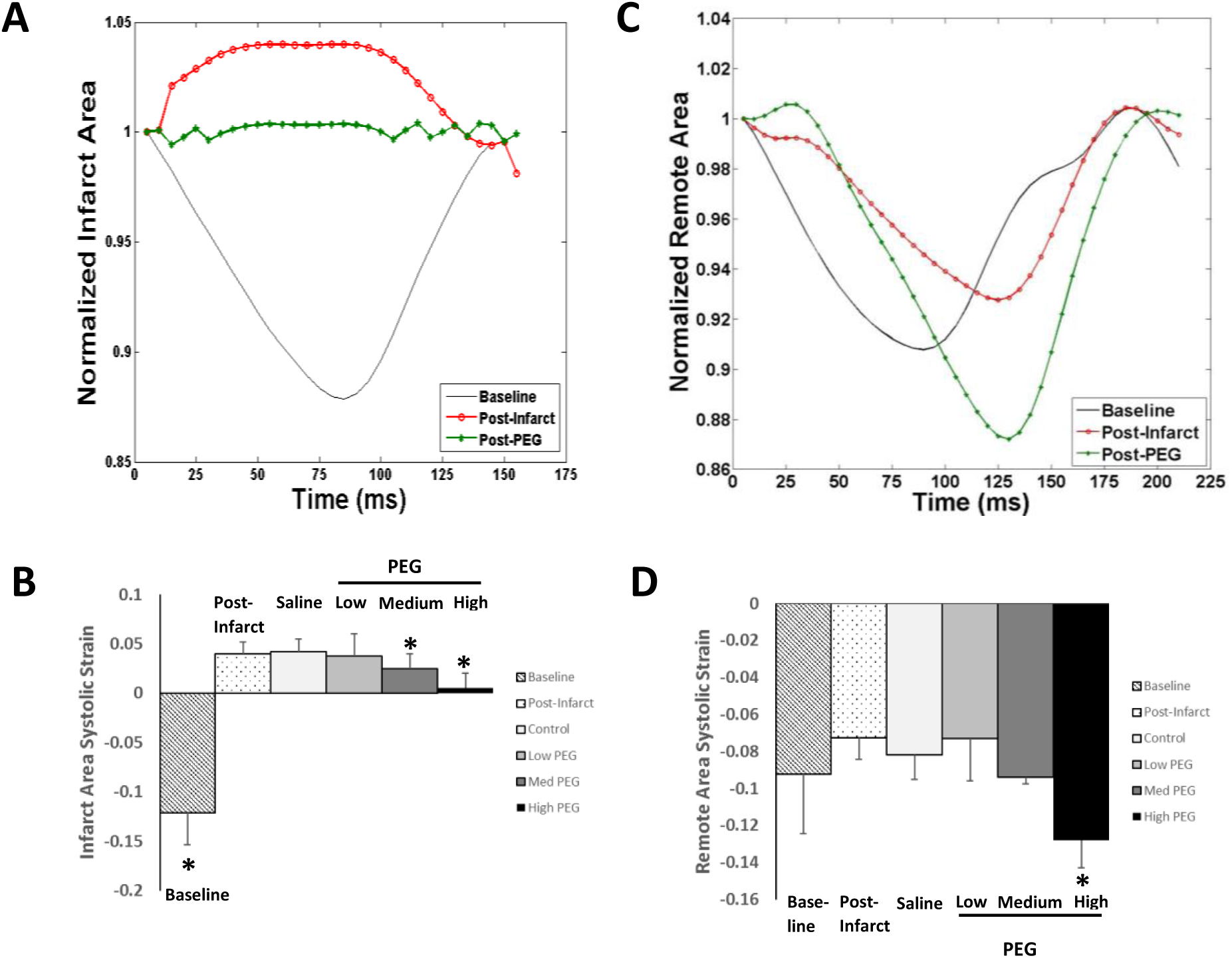
Regional Mechanics. A) Representative normalized infarct area strain curves at baseline, post-infarction, and post-high PEG injection. B) Peak infarct area strain for baseline, post-infarct, control (saline) and 3 PEG doses. Infarct area strain is reduced for medium and high PEG. C) Representative normalized remote area strain curves at baseline, post-infarction, and post-high PEG injection. D) Peak remote area strain for baseline, post-infarct, control, and 3 PEG doses. n=5 subjects/group; *p<0.05 relative to post-infarct.

### Regional Strain

Infarct regional strain became dyskinetic following LAD ligation, which improved following PEG injection. **Figure 4a** illustrates a representative cardiac cycle of systolic strain of the infarct area at baseline, post-infarction, and post-supraphysiologic stiffness (250kPA) PEG gel injection. The dyskinetic region is rendered nearly akinetic. **Figure 4b** shows the peak systolic strains for each mechanical dose of PEG. There was no change in peak systolic strain with control or low stiffness (5kPA) PEG from pre-injection/post-infarct to post-injection. Systolic strain was reduced with medium stiffness (0.025±0.0034 vs 0.0399±0.012; p<0.05) and supraphysiologic stiffness (0.005±0.015 vs 0.0412±0.018; p<0.05) PEG post injection compared to pre-injection/post-infarct. **Figure 4c** illustrates a representative cardiac cycle of systolic strain of the remote area strain at baseline, post-infarction, and post-supraphysiologic PEG injection. Remote area strain also appeared to be reduced post-infarction, however the peak systolic strain was unchanged (**Figure 4d**). Following injection of supraphysiologic stiffness PEG, remote area peak systolic strain increased (−0.128 ± 0.015 vs -0.0724 ± 0.012;p<0.05) post-injection to pre-injection/post-infarct.

### Chronic LV Remodeling

Infarct reinforcement with high mechanical dose, supraphysiologic high PEG prevented adverse post-infarct ventricular remodeling. **Figure 5** illustrates the change in LV morphology and ejection fraction with saline control, low dose (5kPA) PEG, and high (250kPA) PEG. Left ventricular infarct scar area percent was similar between all groups (11±2% saline vs 10±3% Low PEG and 9±3% high PEG) as measured by MRI. Infarct wall thickness however was increased in PEG injection groups (1.9±0.22mm low PEG and 1.7±0.12mm in high PEG) compared to saline control (0.75±0.21mm). There was no difference in LV end-diastolic volume (EDV) or ejection fraction (EF) between control and low dose PEG. High PEG subjects had decreased LV EDV (561±12 vs 695±39; p<0.05) and greater LV EF (44±6% vs 29±6%; p<0.05) compared to control subjects. Peak diastolic filling rate was unchanged between all groups (6.01± 0.30 saline vs 6.28 ±0.22 low PEG vs 5.94±0.12μl/ms high PEG; p=0.39)

**Figure 5.**
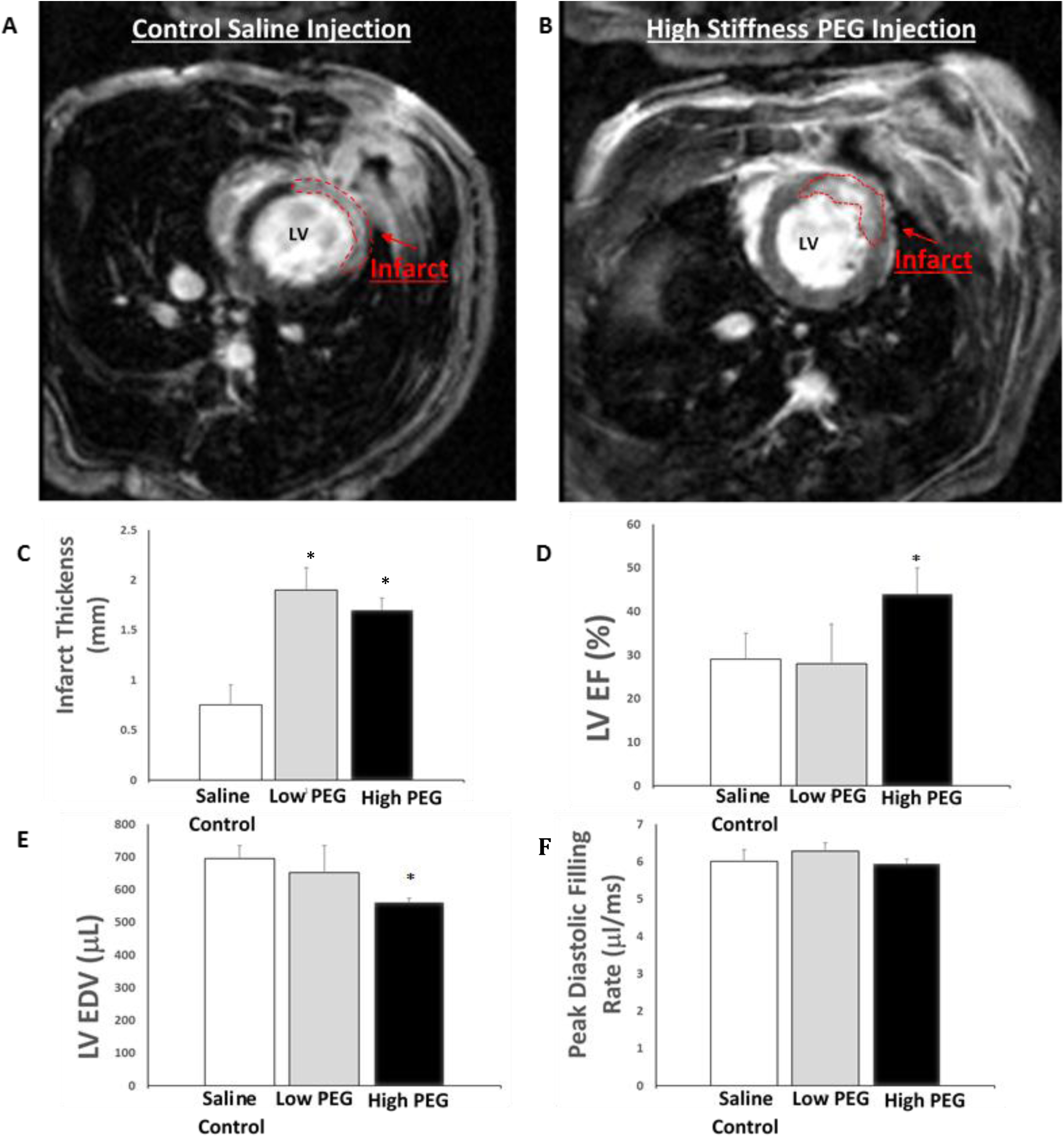
Chronic changes in cardiac morphology and function over 4 weeks. Representative IR-GRE images of infarct for saline (A) and high PEG injections (B). There was no change in infarct size. C) Infarct wall thickness increased with both low and high PEG injection; D Left Ventricular Ejection Fraction (LV EF) increased only with high PEG injection; E) Left Ventricular End-Diastolic Volume (LV EDV) decreased only with high PEG injection. F) There was no change in peak diastolic filling rate. n=3 subjects/group; *p<0.05 relative to saline control.

## Discussion

This study experimentally evaluated the impact of injected intramyocardial biomaterial stiffness on regional and global ventricular mechanics. Infarct reinforcement can eliminate infarct dyskinesia and prevent adverse ventricular remodeling. Importantly, the efficacy of infarct reinforcement is dependent on the mechanical dose of injected biomaterial. Low stiffness biomaterials do not alter regional mechanics and thus, primarily work via augmenting regional wall thickness. Moderate stiffness materials (25kPA) do reduce infarct strain and indices of M_v_O_2_. Supraphysiologic stiff materials (250kPA) reduce infarct area strain and improve remote myocardial strain. At this mechanical dose, materials also acutely increase diastolic stiffness, which may be an important adverse effect, and an upper limit in this model. However, diastolic filling was not significantly impacted chronically indicating this effect may be transient.

Prior studies have demonstrated the potential benefit of infarct reinforcement, however few studies have evaluated ventricular mechanics at different mechanical doses of biomaterial.[15] Wall and colleagues, utilizing finite-element simulations, found that a small fractional change (0.5% to 5.0%) in myocardial wall volume alters mechanics and wall stress.[9] Similarly Kicula and colleagues in *in silico* and finite-element simulations, found that both volume and stiffness of biomaterials influence mechanics.[14] However, *in vivo* only a single biomaterial condition was injected as any increase in stiffness and volume of the wall was found to be beneficial over the injury control. [14] In 2011, the Gorman laboratory at the University of Pennsylvania first demonstrated that intramyocardial injection of tissue filler improved ventricular function in an ovine infarction model. [29] They then subsequently showed that medium stiffness (43kPA) hyaluronic acid had improved efficacy over low stiffness (8kPA) material for chronic remodeling in ovine.[2] A subsequent study demonstrated that medium stiffness gels have a greater reduction of wall stress in finite element analysis of ovine.[30] Based on this seminal work, a wide variety of biomaterials have been studied for infarct reinforcement, however materials with supraphysiologic stiffnesses have not been widely studied. One material, derived from alginate, is currently being investigated in a multi-center randomized control clinical trial involving 78 patients.[31] At 1 year post-injection, infarct reinforcement patients demonstrated improved functional status compared to controls, but no difference was noted in LV remodeling. Rane and colleagues found that wall augmentation alone was insufficient to prevent ventricular remodeling in a rat animal model, which could explain the lack of remodeling noted with soft alginate derived gels.[10] Matsumura and colleagues were the first to demonstrate efficacy with a synthetic biomaterial that was high (257kPA) stiffness.[15] Like Matsumura’s study, we identified an improvement of pump function with supraphysiologic stiffening of the infarct zone. Acute infarct bulging during systole may result in wasted contractile energy by the non-infarcted remote myocardium.[32] Our acute hemodynamic data suggests that reduction of infarct dyskinesia improves remote regional contraction, possibly Taken together, these studies emphasize the importance of biomaterial mechanical properties on infarct reinforcement. Despite the importance of biomaterial stiffness, many studies simply inject a single condition and an optimal condition has not been determined.

The present study provides additional data for rational design criteria for infarct reinforcement biomaterials. An optimized biomaterial may have stiffness properties between 25 and 250kPA. Such a material would reduce infarct area strain and improve remote area strain without increasing diastolic stiffness. In addition to stiffness, gelation time and degradation properties are important factors for delivery and sustainability of treatment.[33, 34] High stiffness materials may be more difficult to deliver to the infarct area in a minimally invasive manner, which is essential to clinical adoption of this therapeutic paradigm. Percutaneous delivery of therapy is ideal and the development of shear-thinning hydrogels allow for easier flow through catheters.[30] Thermosensitive hydrogels, which form hydrogels at body temperature, may also be promising candidates for infarct reinforcement, as they could eliminate the potential lack of cytocompatibility of crosslinking molecules/strategies. [15]

In addition to regional wall augmentation and stiffening, acellular biomaterials may be utilized to engineer the myocardium to allow for myocardial recovery.[35] Gaffey and colleagues have combined sheer thinning hydrogels with endothelial progenitor cells and identified reduced scar formation, improved cell viability, and vasculogenesis.[36] Extracellular matrix derived biomaterials have also been shown to reduce infarct dyskinesia and improve remodeling.[37, 38] Hydrogels may also provide nutrient rich delivery vehicles for cell therapy to induce myocardial recovery and regeneration.[39] Shear-thinning materials, may also protect transplanted cells from mechanical damage experience during syringe injection processes. [40, 41]

This study has several important limitations. The PV analysis and sonomicrometry studies were performed in an acute, open-chested rat under general anesthesia. General anesthesia diminishes compensatory adrenergic and neurohormonal responses, which may augment ventricular function and cardiac output. In this study, we utilized a small animal model, which due to the small heart size allowed placement of sonomicrometry crystals at limited locations on the heart. A more ideal study would place a distinct set of 3-4 crystals in the infarct zone and a separate 3-4 crystals in the remote zone. This is only possible in a large animal model, which would allow space for more crystals. Our chronic study was of limited duration of one month and was performed with 3 subjects per group which may effect study power. Furthermore this study focused on morphometric changes only. The goal of the chronic study was to confirm the findings of our acute study, rather than evaluate the histological and neurohormonal changes that occur with chronic remodeling, which has been demonstrated by others. [15] It is also recognized that the biomaterial stiffness range identified here was in rats and different mechanical properties may ultimately be optimal in humans.

In conclusion, this study demonstrates that infarct reinforcement with acellular biomaterials is a promising therapy to prevent post-infarct ventricular remodeling. The mechanical dose of the biomaterial is an important factor for efficacy. Low stiffness injectable biomaterials do not affect post-infarct regional mechanics. High stiffness supraphysiologic materials eliminate infarct strain and improve remote myocardial mechanics. Given the importance of biomaterial stiffness on remodeling, future studies should identify optimal stiffness by evaluating multiple stiffness conditions. Injectable biomaterials with mechanical properties that exceed the stiffness of native myocardium may maximize efficacy of therapy.

## Data Availability

The raw/processed data required to reproduce these findings cannot be shared at this time due to technical or time limitations.

## Acknowledgements

This study was supported by the Thoracic Surgery Foundation, the University of Virginia Biotechnology Training Grant and the Metro Washington Chapter of ARCS, and the Department of Chemical Engineering, University of Virginia and Translational Health Research Institute of Virginia.

